# Tranquillyzer: A Flexible Neural Network Framework for Structural Annotation and Demultiplexing of Long-Read Transcriptomes

**DOI:** 10.1101/2025.07.25.666829

**Authors:** Ayush Semwal, Jacob Morrison, Ian Beddows, Theron Palmer, Mary F. Majewski, H. Josh Jang, Benjamin K. Johnson, Hui Shen

## Abstract

Long-read single-cell RNA sequencing using platforms such as Oxford Nanopore Technologies (ONT) enables full-length transcriptome profiling at single-cell resolution. However, high sequencing error rates, diverse library architectures, and increasing dataset scale introduce major challenges for accurately identifying cell barcodes (CBCs) and unique molecular identifiers (UMIs) - key prerequisites for reliable demultiplexing and deduplication, respectively. Existing pipelines rely on hard-coded heuristics or local transition rules that cannot fully capture this broader structural context and often fail to robustly interpret reads with indel-induced shifts, truncated segments, or non-canonical element ordering.

We introduce *Tranquillyzer* (TRANscript QUantification In Long reads-anaLYZER), a flexible, architecture-aware deep learning framework for processing long-read single-cell RNA-seq data. *Tranquillyzer* employs a hybrid neural network architecture and a global, context-aware design, and enables precise identification of structural elements - even when elements are shifted, partially degraded, or repeated due to sequencing noise or library construction variability. In addition to supporting established single-cell protocols, *Tranquillyzer* accommodates custom library formats through rapid, one-time model training on user-defined label schemas, typically completed within a few hours on standard GPUs. Additional features such as scalability across large datasets and comprehensive visualization capabilities further position *Tranquillyzer* as a flexible and scalable framework solution for processing long-read single-cell transcriptomic datasets.

## Introduction

Single-cell RNA sequencing (scRNA-seq) has revolutionized our understanding of cellular heterogeneity, enabling the dissection of complex tissues and the identification of distinct cell types and states at unprecedented resolution [1,2]. Traditional scRNA-seq methods, predominantly based on short-read sequencing platforms like Illumina, have facilitated high-throughput gene expression profiling. However, these approaches often fall short in capturing full-length transcripts, limiting insights into isoform diversity, alternative splicing events, and allele-specific expression [3].

The advent of long-read sequencing technologies, such as those offered by Oxford Nanopore Technologies (ONT) and Pacific Biosciences (PacBio), has opened new avenues for scRNA-seq by enabling the sequencing of full-length transcripts. This capability allows for a more comprehensive understanding of transcriptomic complexity, including the identification of novel isoforms and the resolution of complex splicing patterns [3,4]. Despite these advantages, long-read scRNA-seq presents unique computational challenges. High sequencing error rates, diverse and evolving library architectures, and the sheer volume of data generated necessitate robust and adaptable analytical tools [4].

A critical component of scRNA-seq data analysis is the accurate extraction of cell barcodes (CBCs), Unique Dual Indexes (UDIs), and unique molecular identifiers (UMIs), which enable the assignment of reads to their cell of origin and the identification of PCR duplicates, respectively. In long-read sequencing data, high error rates, particularly insertions, deletions, and substitutions, can distort these short sequence motifs, complicating their detection and leading to erroneous cell assignments or quantification [4]. In addition to base-level errors, long-read libraries are also prone to structural artifacts introduced during library preparation, such as incomplete adapter integration, concatemerization, and unintended junctions between fragments [5,6]. These aberrant enzymatic events can obscure the expected read architecture or introduce duplicated elements, further confounding barcode, UDI, and UMI identification. Compounding these issues is the structural variability across library protocols, which differ in the positioning, spacing, and orientation of key elements. Together, these factors present a substantial challenge to the development of generalizable, rule-based parsing strategies for long-read single-cell transcriptomic data.

Existing computational pipelines for barcode, UDI, and UMI extraction often rely on alignment-based methods [7–12]. These methods typically operate under fixed positional assumptions and protocol-specific heuristics, which can break down when insertions, deletions, or truncated elements disrupt expected sequence positions. *Sicelore [12]* relies on the detection of high-purity polyA or polyT stretches and assumes that adapter sequences are located near the termini of reads, making it susceptible to misannotation when read structure deviates from canonical expectations. *scNanoGPS [11]* similarly confines its search for adapter and polyT sequences to the first and last 100 base pairs of each read, a design that can fail in the presence of internal rearrangements or fragmented molecules. *wf-single-cell* (https://github.com/epi2me-labs/wf-single-cell) extracts barcodes and UMIs by aligning the first 100 base pairs of each read to a synthetic reference probe that includes an adapter suffix, polyT, and ambiguous bases representing the barcode and UMI regions. Probabilistic models such as Hidden Markov Models (HMMs)-based strategies [13] are more flexible, but generally model local sequence dependencies and lack the capacity to enforce global structural consistency across the full read. As a result, both approaches may fail to correctly interpret reads with non-canonical element ordering, shifted sequence boundaries, or context-dependent artifacts that are common in long-read protocols.

To address these challenges, we present ***Tranquillyzer*** (TRANscript QUantification In Long reads-anaLYZER), a flexible, architecture-aware deep learning framework specifically designed for processing long-read scRNA-seq data. *Tranquillyzer* employs a hybrid neural network architecture that integrates convolutional neural networks (CNNs) [14] to detect local sequence motifs with bidirectional long short-term memory (BiLSTM) [15] layers to model long-range dependencies across the read. This combination allows the model to simultaneously learn short, discriminative patterns, such as adapter or barcode signatures, and broader contextual relationships between elements, such as their expected order, spacing, and orientation. To further improve sequence labeling accuracy, *Tranquillyzer* supports an alternate model variant that incorporates a conditional random field (CRF) layer [16], which enforces structured transitions between predicted labels and improves consistency in sequence annotation.

## Results

### Tranquillyzer Overview

#### Preprocessing

The feasibility and efficiency of model inference in *Tranquillyzer* are strongly influenced by how reads are retrieved, grouped, and padded - processes that are inherently constrained by available GPU memory. Since it is not practical to load entire single-cell long-read datasets into memory, reads must be processed in manageable chunks. However, neural networks typically require inputs of uniform length, so within each batch, shorter reads must be padded to match the length of the longest read. When batches include highly variable-length reads, particularly those exceeding 10 kb, this padding can lead to substantial memory waste and degraded computational efficiency, underscoring the importance of length-aware batching strategies.

To address this challenge, *Tranquillyzer* implements a length-aware binning strategy that partitions reads into discrete, size-based bins (e.g., 0-499 bp, 500-999 bp, and so on) (Figure 1A). Each bin is written to a separate Parquet file, and binning is performed in parallel across multiple CPU threads to maximize preprocessing throughput. This strategy ensures that reads of similar lengths are grouped together, minimizing unnecessary padding and optimizing GPU memory consumption. In parallel, *Tranquillyzer* generates a lightweight index, mapping each read to its corresponding bin. This index enables rapid retrieval of individual reads for targeted visualization or debugging via the *visualize* sub-command, without reloading the full dataset.

**Figure 1.**
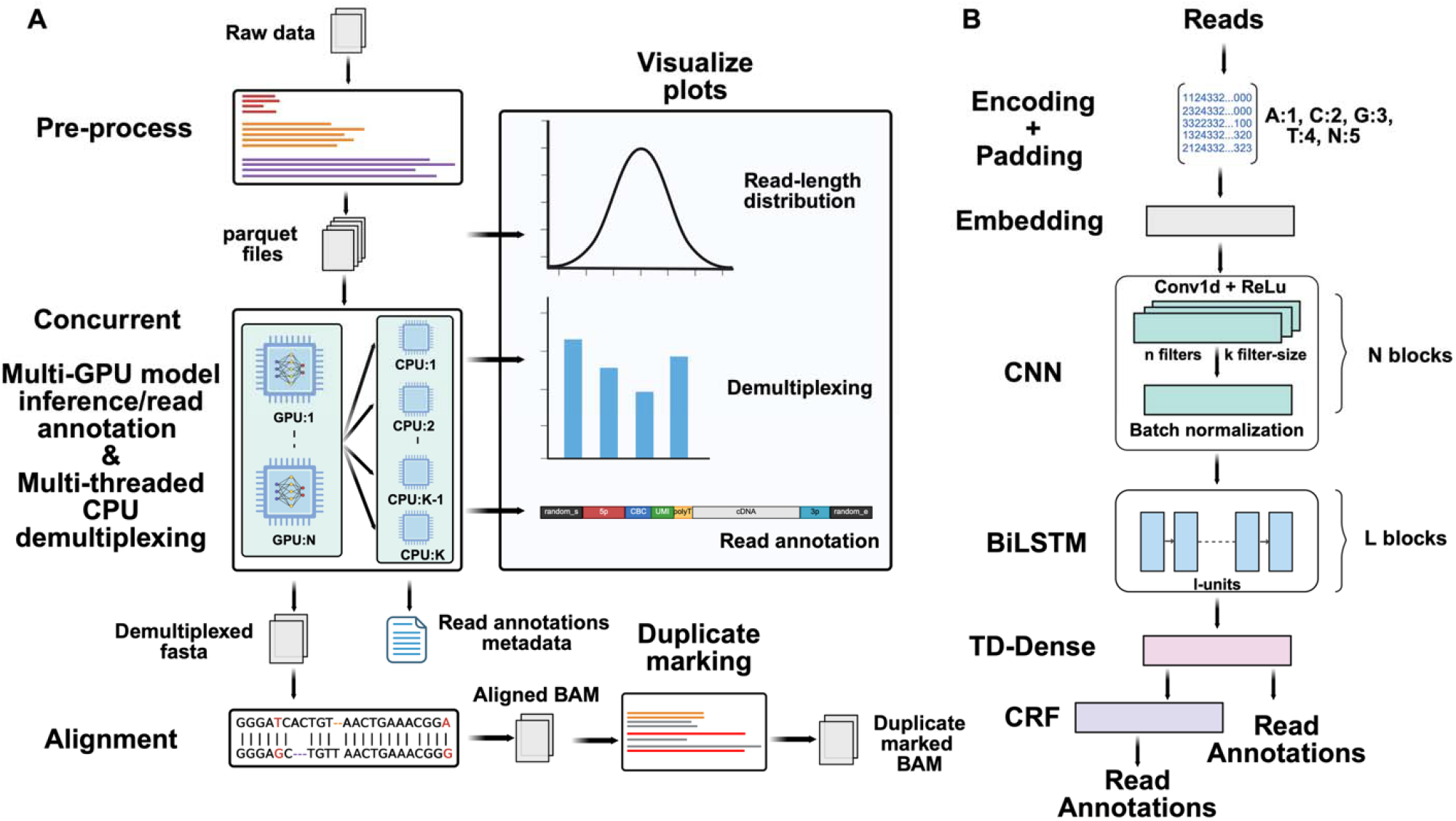
Overview of the *Tranquillyzer* framework. **A)** *Tranquillyzer* accepts raw sequencing data in compressed or uncompressed FASTQ/FASTA format, and first preprocesses them into read-length–based binned Parquet files, for downstream tasks such as annotation, demultiplexing, and visualization (e.g., read-length distributions and per-read annotation visualization). Read annotation and demultiplexing are executed concurrently using multi-GPU inference and multi-threaded CPU processing, respectively. Structural annotations, comprising the start and end coordinates of each feature, inferred cell barcode identity, and filtering status, are stored in annotation metadata files. Successfully demultiplexed reads are separated into a dedicated FASTA file, while ambiguous reads are stored in their own file. Demultiplexed reads are subsequently aligned to a user-defined reference genome using splice-aware alignment, and only primary alignments are retained in the coordinate-sorted BAM output. This file is then used for duplicate marking. **B)** Read annotation is performed using a deep neural network model. Reads are first retrieved from Parquet files in user-defined or default chunks, then encoded and padded to accommodate variable lengths. The model includes a series of convolutional neural network (**CNN**) blocks, each composed of 1D convolutions (**Conv1D**) with **ReLU** activation and optional batch normalization, stacked **N** times as needed for optimal performance. This is followed by a bidirectional long short-term memory (**BiLSTM**) module repeated over **L** layers, a time-distributed dense (**TD-Dense**) layer, and optionally a conditional random field (**CRF**) layer to enforce consistency in base-wise label prediction. The final output is a sequence of per-base annotations identifying structural elements.

#### Neural Network Architecture

*Tranquillyzer* employs a flexible deep learning framework for base-resolution sequence labeling of long-read single-cell RNA-seq reads. Implemented using TensorFlow (v2.15.0) [17], this architecture serves as a generic template from which multiple models can be trained, each tailored to a specific library design or read structure. These models are trained to classify each nucleotide into structural categories such as adapters, primers, polyA/polyT tails, cell barcodes (CBCs), unique molecular identifiers (UMIs), and RNA/cDNA inserts.

Input sequences are first integer-encoded using an ASCII-indexed lookup table, mapping A, C, G, T, and N to values 1 through 5. This encoding step is compiled with *Numba*’s just-in-time (JIT) compiler for fast execution at scale [18]. Reads are padded to the maximum sequence length within each batch, yielding input tensors of shape *(batch_size, max_seq_length)*. An embedding layer maps these integer tokens to dense vector representations, which are then processed by a configurable stack of one or more 1D convolutional layers to detect short-range sequence motifs (e.g., adapter or primer boundaries). This is followed by one or more bidirectional long short-term memory (BiLSTM) layers that capture long-range dependencies and impose global context (**Figure 1B**).

The output layer supports two configurations. In the default variant, a time-distributed dense layer with softmax activation generates a per-base probability distribution over label classes. The softmax function transforms raw scores (logits) at each position into normalized probabilities that sum to one, facilitating probabilistic interpretation and enabling class assignment based on maximum likelihood. Alternatively, a conditional random field (CRF) layer may be used to model structured label transitions jointly across the sequence, improving annotation consistency in noisy or ambiguous regions. The number of layers and architectural parameters, such as filter count, kernel size, LSTM units, and dropout rates, are configurable and tuned for each library-specific model. In practice, a single BiLSTM layer and optional CRF layer have proven effective across a variety of protocols (**Figure 1B**).

Each model is trained using supervised learning on synthetic reads generated by *Tranquillyzer*’s in-house simulation engine. This simulator constructs ground-truth sequences by assembling known adapters and primers with randomly generated CBCs, UMIs, and transcript inserts according to the structural layout defined in a tab-delimited text file. To mimic the characteristic error profile of Oxford Nanopore Technologies (ONT), errors are introduced at user-defined rates: 5% substitutions, 5% insertions, and 6.13% deletions by default. These error-introduced reads are paired with base-level ground truth labels, allowing the model to learn robust annotation despite noise and structural variability. Models with softmax output are trained using categorical cross-entropy loss, while CRF models use a negative log-likelihood objective. Optimization is performed using Adam [19], with learning rate, batch size, and regularization selected per training task.

#### Read Annotation and Demultiplexing

Reads are processed in batches organized by length, as defined during preprocessing. Within each bin, the batch size for annotation inference is dynamically scaled based on the average read length to balance memory usage and throughput. Once batched and encoded, reads are passed through the trained model to infer base-wise label sequences, enabling identification of key structural components such as adapters, cell barcodes (CBCs), unique molecular identifiers (UMIs), cDNA regions, and polyA/T tails.

Model inference is distributed across all available GPU cores using *TensorFlow’s* MirroredStrategy, enabling each batch to be processed concurrently across devices (Figure 1A, B). For each read, the model outputs a probability distribution over labels at every base position, representing the likelihood of different structural elements (e.g., adapter, barcode, UMI, etc.) along the sequence. These probabilities are then decoded into discrete label sequences using a label binarizer corresponding to the trained model. As each batch completes inference, its predictions are offloaded to a pool of CPU threads, configured via a user-defined threads parameter, for postprocessing. This stage includes label decoding, structural validation, barcode correction, and demultiplexing. By overlapping GPU-based inference with CPU-based postprocessing, *Tranquillyzer* maximizes throughput and ensures scalability to large datasets.

From the per-base annotations, contiguous regions are aggregated to identify structural components within each read, including adapters, CBCs, polyA/polyT tails, UMIs, and RNA inserts. The structural validity of each read is assessed by comparing the predicted element order against a protocol-specific label sequence defined in the tab-delimited text file. Reads that conform to the expected structure are marked as valid; those that do not are flagged as invalid.

For structurally valid reads, annotated barcodes are compared against a provided whitelist using the Levenshtein edit distance (**Figure 1A**). Reads with a unique match within a user-defined threshold (default: ≤ 2) are assigned the corresponding barcode, while those that fail to match or yield multiple equally close matches are labeled as ambiguous. Metadata for each annotated read, including the coordinates of detected elements, barcode correction details, and distance metrics, is saved to separate output files: annotations_valid.parquet for valid reads and annotations_invalid.parquet for invalid reads. These files are suitable for both downstream analysis and diagnostic inspection.

In parallel, RNA insert sequences from structurally valid reads with successfully assigned barcodes are written to a demuxed.fasta file, with the corrected CBC embedded in the FASTA header for compatibility with downstream alignment and quantification pipelines. Reads with a valid structural layout but no confidently assigned barcode are instead saved to an ambiguous.fasta file for further inspection or potential rescue.

#### Duplicate Marking

Identifying PCR duplicates using UMIs in large-scale datasets can be computationally intensive, especially if it requires exhaustive pairwise comparisons. To address this challenge, *Tranquillyzer* implements a resource-efficient strategy that significantly narrows the search space. Specifically, it restricts duplicate detection to demultiplexed reads that map to genomic start and end coordinates within a 10 bp window. Within this subset, pairwise Levenshtein distances between UMIs is computed to assess sequence similarity and flag putative duplicates.

The demultiplexed reads are first aligned to a user-specified reference genome using *minimap2* with spliced alignment settings [19] and unmapped reads are discarded before storing into a coordinate-sorted BAM file. *Tranquillyzer* leverages multi-core parallelization to perform duplicate marking independently across reference genome regions or chromosomes (**Figure 1A**).

Aligned reads are loaded into Python via the *pysam* library [20–22]. The tool processes each chromosome or contig separately using user-defined CPU cores. Within each region, reads with identical strand orientation and cell barcode and similar start-end genomic positions are compared for UMI similarity. These filters are configurable, allowing users to relax constraints depending on their experimental design or tolerance for false positives. Duplicates are annotated by setting standard SAM tags and flags for each identified read. Finally, BAM files generated per genomic region are merged to produce a fully duplicate-marked output file, which can be post-processed using tools such as *samtools* [22] for downstream filtering and analysis.

#### Visualization

*Tranquillyzer* incorporates a visualization module to facilitate both quality control and interpretability of annotated reads. It generates several diagnostic plots using the *matplotlib* library [23], including read length distribution histograms to assess sequencing profiles and barcode distance plots that summarize the number of corrected cell barcodes at varying minimum Levenshtein distances from the whitelist. Most notably, *Tranquillyzer* offers detailed, color-coded visualizations of read annotations, where individual structural elements, such as adapter sequences, polyA/T tails, cell barcodes, UMIs, and cDNA regions, are distinctly labeled to enable intuitive exploration of per-read architecture. These visualization features are designed with flexibility in mind and will continue to evolve to accommodate emerging user needs and enhance downstream interpretability (**Figure 1A**).

### Tranquillyzer Accurately Resolves Complex Read Architectures with High Base-Level Accuracy

To assess *Tranquillyzer’s* ability to resolve valid read architectures, we benchmarked its performance against existing tools using simulated single-cell long-read datasets with explicitly defined ground truth. These datasets varied systematically in both read count (ranging from 5 million to 100 million reads, with cDNA lengths between 100-500 bp) and in molecule length (ranging from 500 bp to 2,500 bp, each with 10 million reads), thereby capturing key axes of complexity relevant to long-read single-cell sequencing.

We evaluated *Tranquillyzer* in two annotation modes: the default hybrid mode (*Tranquillyzer_HYB*), which first applies a fast CNN-LSTM model and selectively reprocesses ambiguous reads using a CRF-enhanced model; and the fully CRF-based mode (*Tranquillyzer_CRF*), which applies the more expressive model to all reads. Both modes consistently achieved >99.7% accuracy in structural filtering and >91% demultiplexing efficiency across the full range of dataset conditions (**Figure 2A-C, Supplementary Figure S1A**. These high-performance levels were maintained with increasing sequencing depth and molecule length, indicating robust generalizability to both larger datasets and structurally complex molecules.

**Figure 2.**
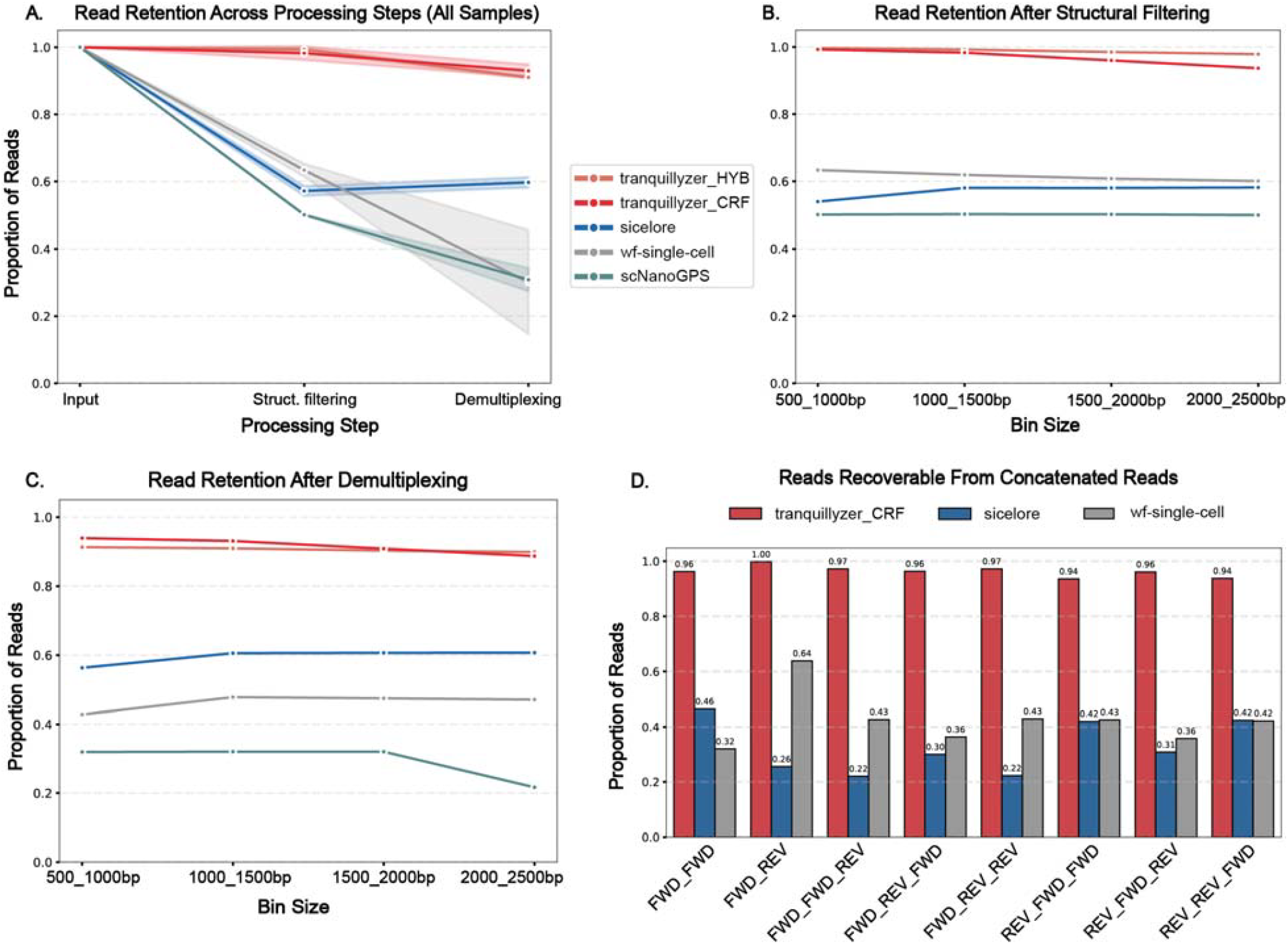
*Tranquillyzer* outperforms existing tools in recovering valid reads and resolving structural artifacts in simulated long-read single-cell datasets. **A)** Read retention across key processing steps - input, structural filtering, and demultiplexing for variable read counts (5-100 million reads) with 100–500 bp cDNA lengths. **B)** Structural filtering efficiency across increasing read lengths (500-2,500 bp with 10 million reads each), with a constant read count of 10M reads. **C)** Demultiplexing performance after structural filtering, stratified by molecule length. **D)** Proportion of sub-fragments recoverable from concatenated reads with known architectures (e.g., FWD_FWD, FWD_REV_FWD, FWD_REV_REV). *scNanoGPS* was excluded from this analysis as it lacks explicit modeling of internal structural complexity.

In contrast, *Sicelore, scNanoGPS*, and *wf-single-cell* showed substantially lower recovery (< 65%) of valid reads. This discrepancy likely arises from their reliance on rigid, motif-centric heuristics. For example, *Sicelore* requires ≥15 nt polyA/T tails with ≥75% purity near read termini; *scNanoGPS* restricts motif detection to the first and last 100 bp of each read; and wf-single-cell enforces strict positional constraints for adapters to classify reads as full-length. These design choices limited their ability to handle noncanonical configurations such as truncated polyA/T regions, internally displaced motifs, or random flanking bases - features deliberately included in the simulated data. While the exact failure mechanisms remain to be fully elucidated, *Tranquillyzer’s* consistent performance across such edge cases underscores its tolerance to structural variation and its strong generalizability.

Interestingly, *Sicelore* exhibited an increase in read retention from the structural filtering to the demultiplexing stage, a pattern not observed with the other tools. This apparent gain reflects *Sicelore’s* retention of reads lacking either a 3′ or TSO motif. To ensure an equitable comparison focused on structurally complete molecules, we explicitly excluded such incomplete reads from *Sicelore’s* structural filtering statistics. In contrast, both *Tranquillyzer* and *wf-single-cell* apply stricter structural criteria upfront ensuring retention of only full-length fragments.

We further evaluated each tool’s ability to identify structural artifacts by focusing on concatenated reads - noncanonical reads formed by the junction of multiple library fragments within a single sequencing read. These chimeric constructs often present valid-looking adapter configurations at the termini while harboring internal structural anomalies that complicate demultiplexing. While tools such as *Sicelore* and *wf-single-cell* incorporate heuristics designed to identify and split such reads, *scNanoGPS* does not explicitly model this class of artifacts and instead relies on fixed-length scanning of read ends. Owing to this architectural limitation in resolving internal complexity, *scNanoGPS* was excluded from this portion of the analysis.

To quantify performances, we curated a dataset of concatenated reads with known architectures, ranging from simple duplications (e.g., FWD_FWD) to more complex patterns (e.g., FWD_REV_FWD) (**Table 1**). For each tool, we measured the proportion of sub-fragments that could be correctly identified, either through explicit read splitting or through structural classification that would enable theoretical recovery, normalized to the known ground truth for each pattern. Given the concordant performance of *Tranquillyzer_HYB* and *Tranquillyzer_CRF* on canonical reads, we conducted this analysis using only *Tranquillyzer_CRF* for simplicity. *Tranquillyzer_CRF* reliably identified these multi-fragment reads across all configurations, achieving theoretical recovery rates exceeding 94% (**Figure 2D**). In contrast, *Sicelore* and *wf-single-cell* recovered only a subset of sub-fragments, likely reflecting a combination of limited sensitivity to internal motif arrangements and the same heuristic constraints that affected their performance on canonical reads.

**Table 1.** Simulated artifactual read types and their descriptions

| Artifact Type | Description |
| --- | --- |
| FWD_FWD | Two forward-oriented fragments joined end-to-end. |
| FWD_REV | A forward-oriented fragment followed by a reverse complement fragment. |
| FWD_FWD_REV | Two forward fragments followed by a reverse complement fragment. |
| FWD_REV_FWD | A forward fragment, then a reverse complement, followed by another forward. |
| FWD_REV_REV | A forward fragment followed by two reverse complement fragments. |
| REV_FWD_FWD | A reverse complement fragment followed by two forward fragments. |
| REV_FWD_REV | A reverse complement fragment followed by a forward, ending with another reverse complement. |
| REV_REV_FWD | Two reverse complement fragments followed by a forward fragment. |

These results establish *Tranquillyzer* as high-fidelity tools for resolving complex read architectures in long-read single-cell data. Their near-perfect classification of both canonical and artifactual reads across a broad range of sequencing conditions supports their robust applicability to real-world datasets.

### Tranquilizer Effectively Resolves Complex Read Artifacts in Real Single-Cell Long-Read Transcriptomes

To assess performance in the context of real biological data, we applied all four tools to a publicly available single-cell long-read transcriptome as described in methods. In the absence of ground truth, we utilized *Tranquillyzer_CRF*-derived structural annotations as a consistent reference framework to evaluate the nature and integrity of recovered reads (**Figure 3A**).

**Figure 3.**
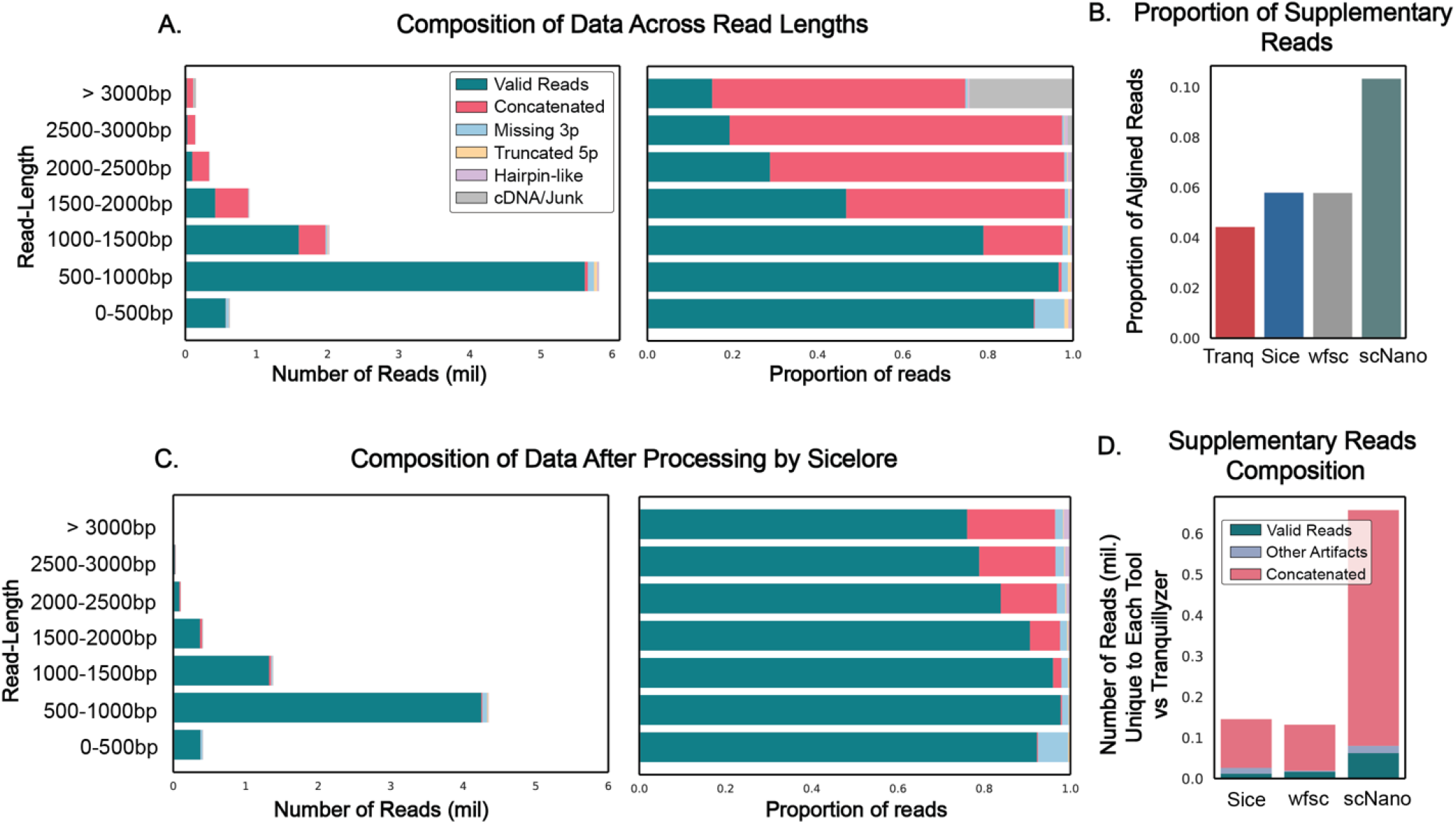
Tranquillyzer provides high-fidelity structural classification and resolves complex artifacts in real single-cell long-read transcriptome datasets. **A)** Read composition stratified by length bin in a real dataset processed with *Tranquillyzer_CRF*. Left panel shows absolute read counts per bin; right panel displays the relative proportions of read categories, including valid single fragments, concatenated molecules, and various classes of truncated or artifactual reads. **B)** Proportion of mapped reads containing supplementary alignments across tools. **C)** Read composition across read-length bins after processing with *Sicelore*, with subsequent structural reclassification using *Tranquillyzer_CRF*. **D)** Composition of supplementary alignments unique to each tool (i.e., not shared with *Tranquillyzer_CRF*).

*Tranquillyzer* demonstrated the highest total yield of theoretically recoverable full-length molecules (**Supplementary Figure S2A**), substantially exceeding the other tools. This elevated recovery reflects its unique capacity to both accurately identify single-fragment canonical molecules and detect multi-fragment concatenated reads. The ability to resolve complex read architectures contributes to a more comprehensive and structurally faithful representation of the transcriptome.

*scNanoGPS* produced the highest proportion of reads with supplementary alignments (>10%) among all evaluated tools, consistent with its limiting motif detection to fixed-length regions at the read termini. This design inherently constrains its ability to resolve internal structural complexity, such as concatenated sub-fragments, thereby increasing the likelihood of misclassifying chimeric molecules as valid single fragment reads. In contrast, *Sicelore* and *wf-single-cell* yielded fewer supplementary alignments. However, a focused examination of supplementary reads unique to these tools (i.e., those not overlapping *Tranquillyzer* supplementary reads), revealed that the vast majority (>80%) originated from molecules annotated by *Tranquillyzer* as concatenated (**Figure 3D, Supplementary Figure S2B**). This suggests that, despite some ability to detect or partially resolve concatenated constructs, both tools continue to misclassify a fraction of structurally artifactual molecules as valid, potentially confounding downstream analyses. These results highlight the need for precise structural delineation in long-read transcriptomic workflows and demonstrate *Tranquillyzer’s* ability to identify and characterize complex artifacts with high resolution.

Notably, since *Sicelore* retains adapter sequences in output reads, we reanalyzed its processed data using *Tranquillyzer* to assess residual artifact content. This secondary classification confirmed a marked reduction in concatenated molecules (**Figure 3C**), indicating partial efficacy of *Sicelore’s* splitting strategy. However, residual artifacts, particularly in longer reads, persisted. This trend was consistent with observations from raw data (**Figure 3A**), wherein the prevalence of artifactual reads increased with read length. Although the dataset analyzed had a median read length of 855 bp (**Supplementary Figure S2C**), this pattern is expected to be further exacerbated in datasets generated using protocols optimized for longer transcripts.

These findings demonstrate that while existing tools may achieve superficially comparable full-length yields (**Figure 3B, Supplementary Figure S2A**), *Tranquillyzer* uniquely couples high recovery rates with rigorous structural fidelity. Its capacity to identify both canonical and artifactual molecules positions it as a robust solution for high-resolution annotation in long-read single-cell transcriptomics.

### Read Annotation Visualization Enables Dissection of Structural Artifacts

To complement the structural annotation and quantitative benchmarking presented above, we implemented read-level visualization capabilities in *Tranquillyzer* to support manual inspection of complex molecules. These visualizations serve a critical interpretive function by enabling users to distinguish true biological features from sequencing artifacts that might otherwise confound downstream analyses.

As a representative example, we examined a read that exhibited supplementary alignments in the outputs of *Sicelore, wf-single-cell, and scNanoGPS* (**Figure 4**). All three tools aligned the 5′-anchored cDNA segment as the primary alignment, while the downstream cDNA region, originating from a concatenated subfragment, was assigned a supplementary alignment. Importantly, *Tranquillyzer* annotated this molecule as a double-fragment concatenated read (concatenated reads x2 (+:2), indicating two forward-oriented sub-fragments within the read) and flagged it as structurally artifactual. This case illustrates how, in the absence of accurate structural dissection, chimeric molecules may be misinterpreted as biologically meaningful features such as fusion transcripts or structural variants. Indeed, the presence of a split alignment pattern, primary followed by supplementary, could easily be mistaken for a legitimate genomic fusion event when in fact it reflects an artifact of library preparation or sequencing.

**Figure 4.**
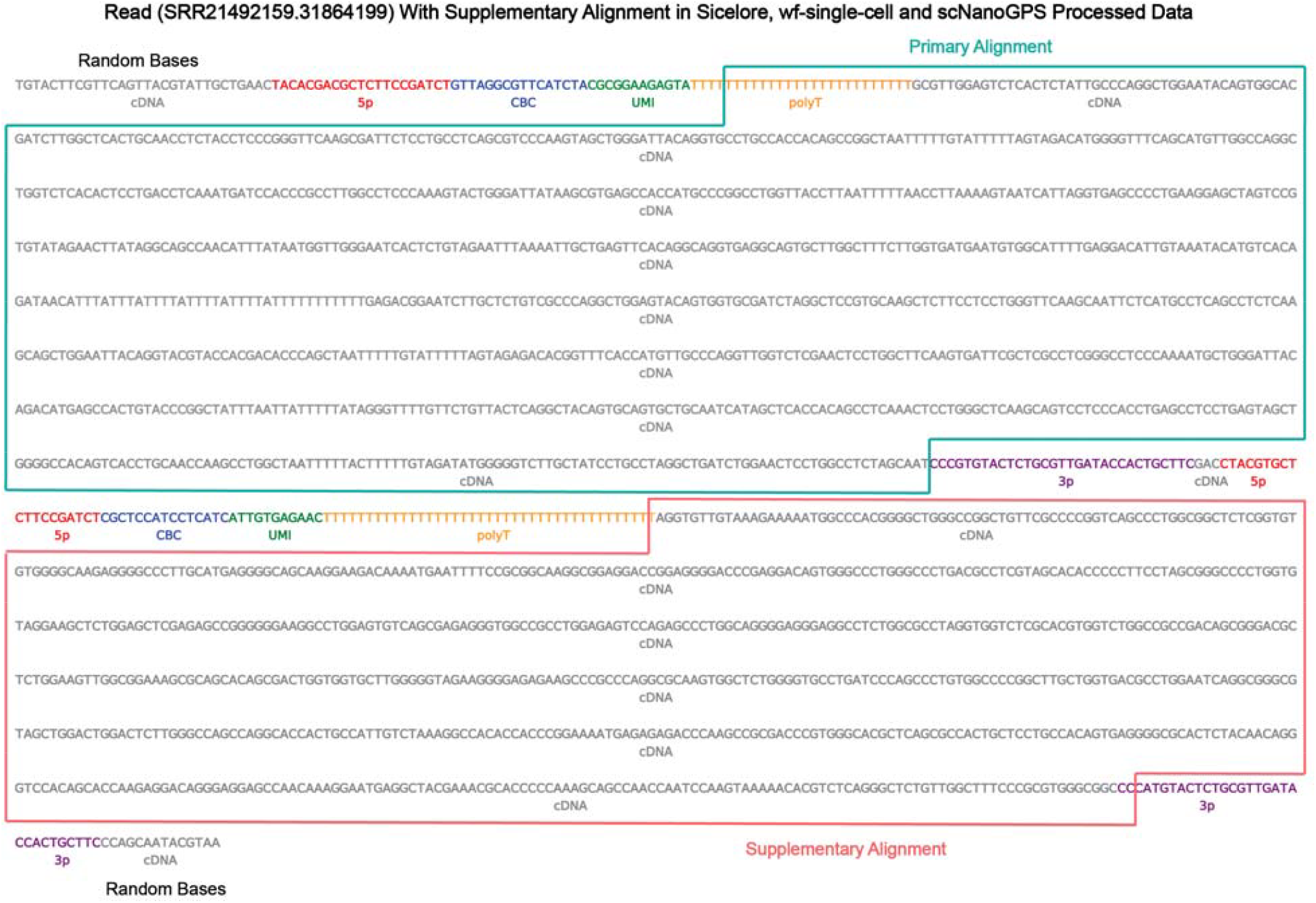
Read annotation visualization reveals misclassified structural artifacts across other tools. A representative read (SRR21492159.31864199) exhibiting supplementary alignments in the outputs of *Sicelore, wf-single-cell, and scNanoGPS* is shown. All three tools aligned the upstream cDNA fragment as the primary alignment (green box), while the downstream fragment - originating from a concatenated fragment - was misclassified as a valid supplementary alignment (red box). In contrast, *Tranquillyzer* annotated this read as a concatenated read composed of two distinct sub-fragments, flagging it as structurally artifactual. Color-coded labels denote annotated elements including adapters (5′, 3′), polyT tail, cell barcode (CBC), UMI, and cDNA.

The ability to visualize such reads with base-level label overlays, including adapter motifs, cell barcodes, UMIs, polyT tails, and cDNA segments, allows users to directly verify structural integrity and resolve ambiguity in interpretation. This diagnostic resolution is particularly valuable in single-cell contexts, where individual reads carry amplified significance, and artifacts may propagate false transcript structures or splicing events.

In summary, *Tranquillyzer’s* integrated visualization module enhances the interpretability and trustworthiness of annotated long-read datasets. By combining high-fidelity structural classification with intuitive visual diagnostics, it provides a comprehensive framework for ensuring both the quality and biological plausibility of transcriptomic inferences in single-cell long-read sequencing.

### Tranquillyzer Demonstrates Efficient Scalability Across Read Depths and Lengths

To assess scalability under realistic sequencing demands, we profiled the runtime performance of *Tranquillyzer* against existing pipelines across two orthogonal experimental conditions: increasing read counts with fixed read length (0-500bp bin) and increasing read length with a constant number of reads (10 million per bin) (**Supplementary Figure S1B-C**). All pipelines were executed using 64 CPU cores. Additionally, *Tranquillyzer* leveraged 4 parallel GPUs for model inference, consistent with its design for high-throughput annotation acceleration.

Direct comparisons of absolute runtime, however, require careful contextualization. The evaluated tools implement key processing steps in varying order and depth, precluding direct comparisons. For instance, both *wf-single-cell* defers barcode correction until after alignment and only applies correction to reads that have already passed structural filtering. In contrast, *Tranquillyzer* performs barcode correction immediately after annotation and structural filtering, prior to alignment. Since previous benchmarking (**Figure 2**) established that *Sicelore, scNanoGPS*, and *wf-single-cell* recover fewer than or near 60% of structurally valid reads in simulated datasets, the effective input for barcode correction and demultiplexing is substantially reduced relative to *Tranquillyzer*, reducing their runtime.

Moreover, downstream analysis steps beyond demultiplexing also differ across pipelines. For example, *wf-single-cell* proceeds to gene-level quantification and UMAP generation, while *Tranquillyzer* currently terminates at duplicate marking. This design choice reflects our focus on *Tranquillyzer’s* core innovations, accurate annotation of structural elements, robust classification of valid and artifactual reads and high-fidelity demultiplexing, rather than on complete transcriptomic quantification, which is supported by mature software packages downstream of our framework.

Despite these differences in pipeline scope and structure, *Tranquillyzer* exhibited strong scalability and competitive performance. Both its CRF-only and hybrid modes scaled linearly with increasing read counts and read lengths (**Supplementary Figure S1B-C**), maintaining practical runtimes even for 100 million reads or reads exceeding 2,000 bp. The hybrid mode (*Tranquillyzer_HYB*) achieved considerable speed gains over the full CRF implementation (*Tranquillyzer_CRF*) while retaining high accuracy (**Figure 2**), offering a favorable balance of speed and performance for most applications. By contrast, *scNanoGPS* exhibited disproportionately longer runtimes, particularly as read lengths increased.

These results highlight that *Tranquillyzer* not only achieves superior structural and demultiplexing accuracy but does so with a level of computational efficiency that supports deployment on large-scale single-cell long-read datasets. Its GPU-accelerated design and modular preprocessing framework position it as a scalable solution for the evolving landscape of long-read transcriptomics.

## Discussion

Long-read transcriptomics offers unprecedented resolution in capturing full-length RNA molecules and transcript isoforms at single-cell resolution [1,2]. However, this potential remains underutilized due to the limitations of existing preprocessing pipelines in accurately parsing structurally complex and heterogeneous reads. In this work, we present *Tranquillyzer*, a novel neural network-based framework that addresses these challenges through high-resolution read annotation, structural filtering, and demultiplexing. While tailored handle single-cell long-read transcriptomics, its architecture and modular design are equally applicable to bulk long-read datasets as well as other sequencing modalities such as whole-genome sequencing (WGS).

*Tranquillyzer* distinguishes itself by its ability to accurately label key structural motifs, including barcodes, UMIs, adapters, and transcript boundaries, at single-base resolution. Unlike heuristic-based tools, it leverages a CNN-LSTM model architecture with an optional CRF layer for context-aware predictions, allowing precise classification even in the presence of noncanonical configurations, shortened motifs, or internal structural artifacts. Our benchmarking on simulated datasets demonstrates that *Tranquillyzer* achieves >99.7% structural annotation accuracy and >91% demultiplexing efficiency, outperforming existing tools such as *Sicelore, wf-single-cell, and scNanoGPS*, which exhibit substantially lower recovery of valid reads due to rigid motif-centric heuristics. While these results underscore *Tranquillizer*’s prediction consistency, the optional CRF layer, may increase runtime slightly compared to simpler architectures, especially in large-scale datasets. Although the hybrid mode mitigates this, users may need to balance between runtime and accuracy based on application.

Critically, *Tranquillyzer* not only identifies canonical single-fragment molecules with high fidelity, it also systematically flags multi-fragment concatenated reads and other structural artifacts. Such artifacts, if unrecognized, can propagate through downstream analyses, inflating rates of supplementary alignments and potentially being misinterpreted as splicing or structural variants. Furthermore, they may confound quantification by including multiple transcripts within a single CBC-UMI combination, leading to overcounting. *Tranquillyzer’s* ability to annotate and classify such reads with granularity is essential for ensuring transcriptomic fidelity and interpretability. Despite its sophisticated annotation framework, *Tranquillyzer* remains computationally efficient. It demonstrates favorable scaling with increasing read counts and read lengths, supported by multi-GPU acceleration and parallelized I/O, achieving competitive runtimes relative to other pipelines.

Complementing its analytic precision, *Tranquillyzer* incorporates interactive read annotation visualization, allowing direct inspection of motif boundaries, structural organization, and classification outcomes. We illustrate the utility of this feature through the analysis of an exemplar read that aligned as both primary and supplementary in all alternative pipelines but was correctly annotated by *Tranquillyzer* as a concatenated artifact. Without such visualization and structural awareness, such molecules might be falsely interpreted as biologically meaningful fusion events.

*Tranquillyzer* is also highly flexible, supporting known library designs such as 10x Genomics 3′ and 5′ scRNA-seq protocols, and is readily extensible to emerging chemistries. It is well-suited to handle complex platforms such as MAS-ISO-seq, where multiple full-length cDNAs are embedded within single reads. The conceptual overlap between splitting concatenated artifacts and demultiplexing MAS-ISO-seq reads enables seamless adaptation of *Tranquillyzer’s* annotation and parsing framework to this emerging library type. Work is currently underway to integrate this functionality into future versions.

While *Tranquillyzer* offers multiple advantages, some limitations warrant discussion. First, although *Tranquillyzer* supports model training for new or custom library types, achieving optimal performance requires a well-trained model tailored to the specific sequencing chemistry. Creating accurately labeled training data for such models can be challenging, particularly for users without prior experience in model development or annotation curation. In these cases, performance may be limited until appropriately labeled examples representative of the target library design are generated. Second, while *Tranquillyzer* was trained and benchmarked on systems equipped with four NVIDIA L40 GPUs, the minimum GPU memory requirements for reliable inference and training remain to be systematically determined. As with many neural network models, users may encounter out-of-memory (OOM) issues depending on hardware configurations, particularly when working with large batches or long reads. Future work will include formal benchmarking across diverse GPU setups to better characterize resource needs. Lastly, while effective for known forms of chimerism and fragmentation, *Tranquillyzer* may miss rare or uncharacterized artifacts not captured during model training. Continued refinement through periodic updates will be essential to maintaining *Tranquillyzer*’s robustness and adaptability across evolving use cases.

In summary, *Tranquillyzer* offers a robust, accurate, and scalable solution for preprocessing single-cell long-read transcriptomic data. By coupling deep learning–based annotation with artifact-aware structural filtering, intuitive visualization, and broad applicability across library types, it represents a significant advance toward trustworthy and reproducible analysis of complex transcriptomes.

## Materials and Methods

### Benchmarking Data

To evaluate *Tranquillyzer*’s performance relative to existing tools, we leveraged simulated datasets where the ground truth was explicitly defined. We selected the 10x Genomics 3’ single-cell RNA-seq (scRNA-seq) library, one of the most widely used formats for transcript quantification using Oxford Nanopore Technologies (ONT), as the basis for both read simulation and benchmarking with real datasets.

Simulated reads were constructed by concatenating library elements in the canonical 10x 3’ sequencing order: 5’ adapter → cell barcode (CBC) → unique molecular identifier (UMI) → polyT stretch → 3’ cDNA fragment. CBCs were randomly sampled from a pool of 6,500 barcodes drawn from the official 10x whitelist of 3 million barcodes. To mimic the variability observed in real sequencing data, random bases (of randomly selected lengths ranging from 0 to 70 bp) were appended to both ends of each read. Sequencing errors were simulated using empirically defined probabilities: substitution rate = 0.05, insertion rate = 0.05, and deletion rate = 0.06. Alongside the simulated FASTQ files, a tab-separated metadata file was generated to track the ground-truth CBC and UMI sequences associated with each read.

Two primary factors influence the computational performance and scalability of long-read processing tools: total number of reads and read length. To systematically assess performance under varying conditions, we generated two series of datasets. In the first series, we increased read counts from 5 million to 100 million reads (5M, 25M, 50M, 75M, and 100M) while maintaining a consistent cDNA insert length distribution between 100 and 500 bp. In the second series, we simulated reads of increasing lengths by expanding the insert size from 500 bp to 2,500 bp in 500 bp increments (i.e., 500-1,000 bp, 1,000-1,500 bp, 1,500-2,000 bp, and 2,000-2,500 bp).

Beyond detecting structurally valid reads and performing demultiplexing, a critical challenge for long-read tools is the accurate identification and flagging of structurally artifactual reads. Given the well-documented prevalence of structural artifacts in long-read scRNA-seq libraries [5,6], we also generated synthetic datasets containing several distinct categories of artifactual reads (**Table 1**). These were used to assess each tool’s ability to correctly classify and discard noncanonical read structures.

To complement the simulation-based assessments and evaluate *Tranquillyzer*’s performance under real-world conditions, we analyzed a publicly available long-read single-cell transcriptomic dataset generated using Oxford Nanopore’s PromethION platform. The dataset was retrieved from the Gene Expression Omnibus (GEO) under accession number GSE212945 (SRA ID: SRR21492159). To facilitate accurate cell barcode assignment and validation, we also downloaded the corresponding barcode whitelist derived from matched short-read sequencing (SRA ID: SRR21492168), as provided in the same study. This allowed direct comparison of *Tranquillyzer*’s demultiplexing marking performance using validated barcoding information from a hybrid short- and long-read experimental design.

### Benchmarking Parameters

To ensure a consistent and equitable performance comparison, all benchmarking analyses were conducted using the same high-performance computing environment. Each tool - Tranquillyzer, *Sicelore, scNanoGPS, and wf-single-cell)* - was executed using 64 CPU cores per run. Additionally, *Tranquillyzer* leveraged four NVIDIA L40S GPUs to accelerate its annotation pipeline, which includes deep learning–based inference. For barcode correction, each method was configured to utilize common whitelists with a Levenshtein-distance threshold of 2, harmonizing error tolerance across tools and ensuring that variation in correction stringency did not bias downstream metrics.

*Tranquillyzer* used the following core subcommands in sequential order: preprocess, annotate-reads, align, and dedup. For all steps, the number of threads was set to *--threads 64* to ensure full utilization of the allocated CPU resources. During annotation, *--chunk-size* was set to 300,000 to optimize GPU memory usage and throughput. For alignment, the *--preset splice* option was applied to enable spliced alignment using minimap2. For duplicate marking, the parameters *--lv-threshold 2* (Levenshtein distance for UMI deduplication) and *--per-cell* (per-cell deduplication) were specified. All other parameters utilized their default settings.

For each of the benchmarking datasets, *Sicelore* version 2.1 was run following a slightly modified version of the provided *Nextflow* pipeline [24]. Three steps were included in the pipeline: 1) scanning the raw reads and extracting valid barcodes, 2) aligning reads passing the scanning filters, and 3) assigning UMIs to the reads. Step 1 was run via: *java -jar Nan oporeBC_UMI_finder-2.1.jar scanfastq -d <FASTQ_DIR> -o passed --ncpu <N_CPUs> -- bcEditDistance 1 --compress --cellRangerBCs <CBC_WHITELIST>*. Individual FASTQs passing the scanning step were merged into a single FASTQ for downstream processing. Valid barcodes were extracted via: *java -jar Sicelore-2.1.jar SelectValidCellBarcode -I BarcodesAssigned.tsv -O BarcodesValidated.csv -MINUMI 1 -ED0ED1RATIO 1*. Reads were aligned with minimap2 (version 2.28) and coordinate-sorted with samtools (version 1.20). The command used was: *minimap2 -ax splice -uf --sam-hit-only -t <N_CPU> --junc-bed <SPLICE_JUNCTION_BED> <REFERENCE> merged.fq.gz* | *samtools view -bS -@ <N_CPU> -* | *samtools sort -m 2G -@ <N_CPU> -o passed.bam - && samtools index passed.bam*. Reads were aligned to the hg38 reference FASTA and the splice junction BED was generated with the *gff2bed* utility in *paftools.js* (provided with minimap2). UMIs were assigned to reads via: *java -jar NanoporeBC_UMI_finder-2.1.jar assignumis --inFileNanopore passed.bam -o passedParsed.bam --annotationsFile <REFFLAT>*. The REFFLAT file was generated using: *gtfToGenePred -genePredExt -geneNameAsName2 <ANNOTATION_GTF> tmp.refFlat* followed by *awk -F”\t” ‘BEGIN{OFS=“\t”} {print $12, $1, $2, $3, $4, $5, $6, $7, $8, $9, $10}’ tmp.refFlat > hg38.refFlat*. For each command, 64 CPUs were used with 256 GB of memory requested and a maximum run time of 48 hours allowed.

scNanoGPS is a python script that generates a shell script which calls individual python scripts that perform barcode detection, barcode collapsing, UMI correction, and demultiplexing. scNanoGPS v2.0 was downloaded from https://github.com/gaolabtools/scNanoGPS. scNanoGPS requires python 3.11 and a conda environment was created with the following packages installed: biopython 1.80 [25], distance 0.1.3 (https://pypi.org/project/Distance/), matplotlib 3.8.2 [23], pandas 2.1.4 [26], pysam 0.23.0 [20], seaborn 0.13.1 [27]. Other necessary dependencies were installed with bioconda: minimap2 2.30-r1287 [19], samtools v1.21 [22], tabix 1.21 [28], spoa v4.1.0 (https://github.com/rvaser/spoa), subread v2.1.1 [29], and longshot v1.0.0 [30]. To run scNanoGPS, *scanner.py* scans reads to identify polyA and the TruSeq Read1 adapter. It then extracts the CB and UMI from reads that have a valid architecture, generating a processed FASTQ file that contains these reads. This script was run with default options except *--min_read_length* was set to 100 (default 200). Next, *assigner.py* collapses cell barcodes using an edge detection strategy. Since v2.0, *assigner.py* also accepts a whitelist which we provided where relevant. This tool was also run with default settings except the minimum read number was set to 1 when using a whitelist to avoid false negatives. The next step is *curator.py* which demultiplexes the processed FASTQ file from *scanner.py* using the corrected cell barcodes from *assigner.py* so that each cell gets a single FASTQ file with all reads; these are then mapped to the reference genome using minimap2 in spliced alignment mode.

wf-single-cell is a nextflow pipeline used for the preprocessing of single-cell or spatial transcriptomics experiments performed using 10x technology. The wf-single-cell workflow first identifies adapters using vsearch [31]. Then wf-single-cell splits concatenated reads based on the presence of multiple sets of these adapters in a single read and assigns adapter configurations to each read. The pipeline then uses parasail [32] to identify UMI and cell barcode sequences in the each read. Reads are mapped to the selected reference genome using minimap2. Barcodes are corrected using the appropriate 10x whitelist and reads are assigned to the appropriate transcript using a stringtie2 [33] generated transcriptome. UMI-tools [34] with gene clustering and a Levenshtein distance of 2 is used to correct and assign UMIs to reads. Finally, the pipeline generates gene and transcript expression matrices and tags bam files with the corrected and uncorrected cell barcode and umi sequences. For each of the benchmarking datasets wf-single-cell version v3.1.0-g4580fc3 was used. The nextflow pipeline was run using the following parameters for nextflow: -c $NEXTFLOW_CONFIG, -w $WORKSPACE, -r v3.1.0-g4580fc3, -profile singularity, -with-trace. The pipeline was run using the following parameters for wf-single-cell: --fastq $FASTQ, --expected_cells 1000, -- matrix_min_cells 0, --matrix_min_genes 0, --kit 3prime:v3, --ref_genome_dir refdata-gex- GRCh38-2024-A, --out_dir $OUT_DIR, --threads 64, --sample simulated_data. For each benchmarking dataset, the maximum memory used was 256 GB.

## Data and Code Availability

*Tranquillyzer* is available for installation at: https://github.com/huishenlab/tranquillyzer.git

All scripts used for benchmarking in this study are accessible at: https://github.com/huishenlab/tranquillyzer_manuscript_code.git

## Acknowledgments

Computation for the work described in this paper was supported by the High-Performance Cluster and Cloud Computing (HPC3) Resource at the Van Andel Research Institute.

