## Supplementary Figures for "Tranquillyzer: A Flexible Neural Network Framework for Structural Annotation and Demultiplexing of Long-Read Transcriptomes"

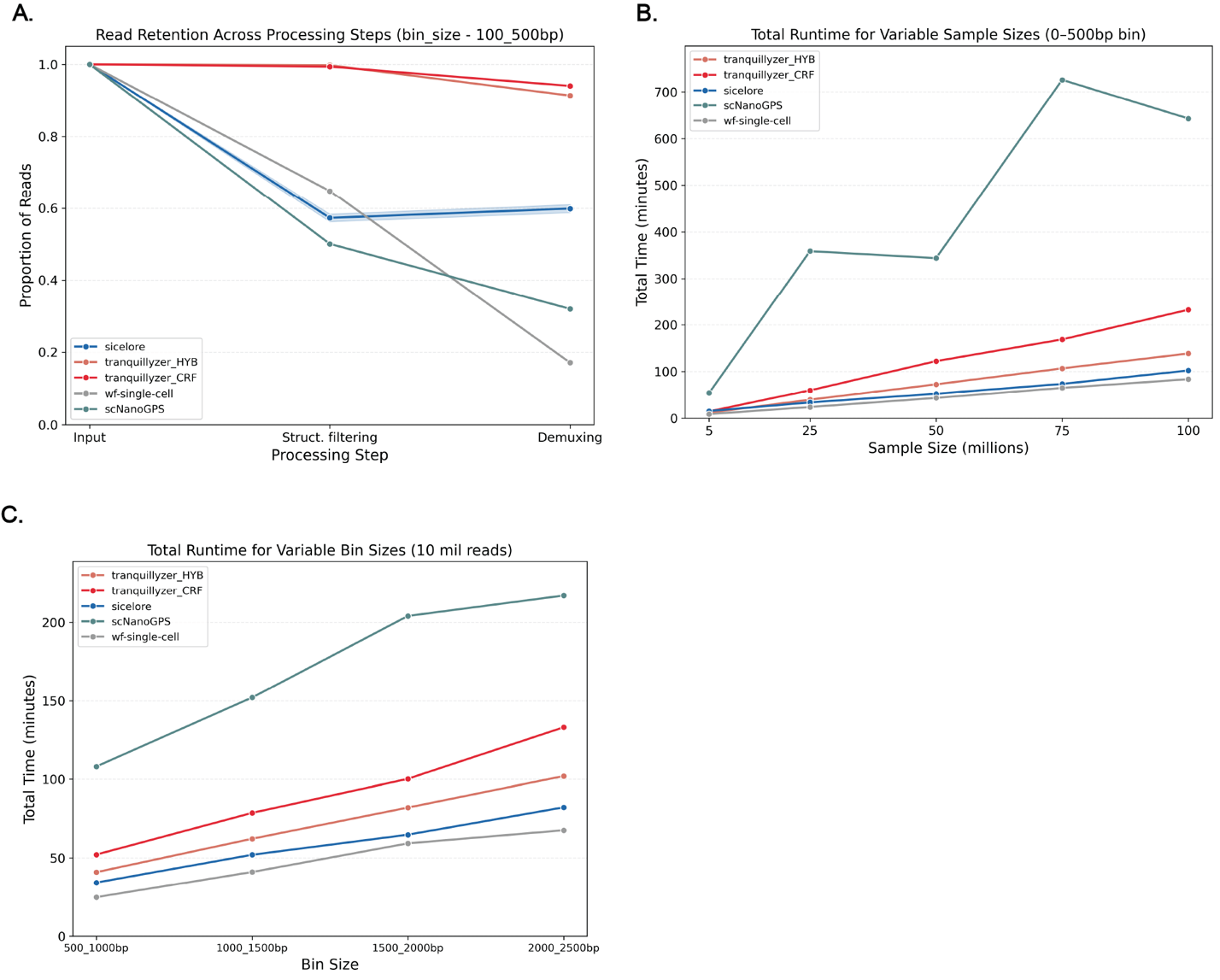

**Supplementary Figure S1: *Tranquillyzer* exhibits robust scalability across increasing sequencing depths and read lengths while maintaining high accuracy and practical runtime.** (A) Total runtime (in minutes) across varying sample sizes (5-100 million reads) within the 0-500 bp bin. *Tranquillyzer_HYB* and *Tranquillyzer_CRF* scale linearly, with the hybrid mode offering substantial speed improvements while retaining high accuracy. (B) Runtime for datasets with fixed sample size (10 million reads) and increasing read lengths (bin sizes ranging from 500 to 2500 bp). Despite increasing molecule complexity, *Tranquillyzer* maintained competitive performance. *scNanoGPS* runtime increased steeply with longer reads, whereas other tools exhibited more modest scaling behavior. All tools were run using 64 CPU cores; *Tranquillyzer* additionally utilized 4 NVIDIA L40S GPUs for model inference. (C) Read retention across processing steps for the 100-500 bp bin (5-100 million reads). *Tranquillyzer* consistently retained the highest proportion of reads through structural filtering and demultiplexing stages, underscoring its superior sensitivity and accuracy in resolving valid molecules.

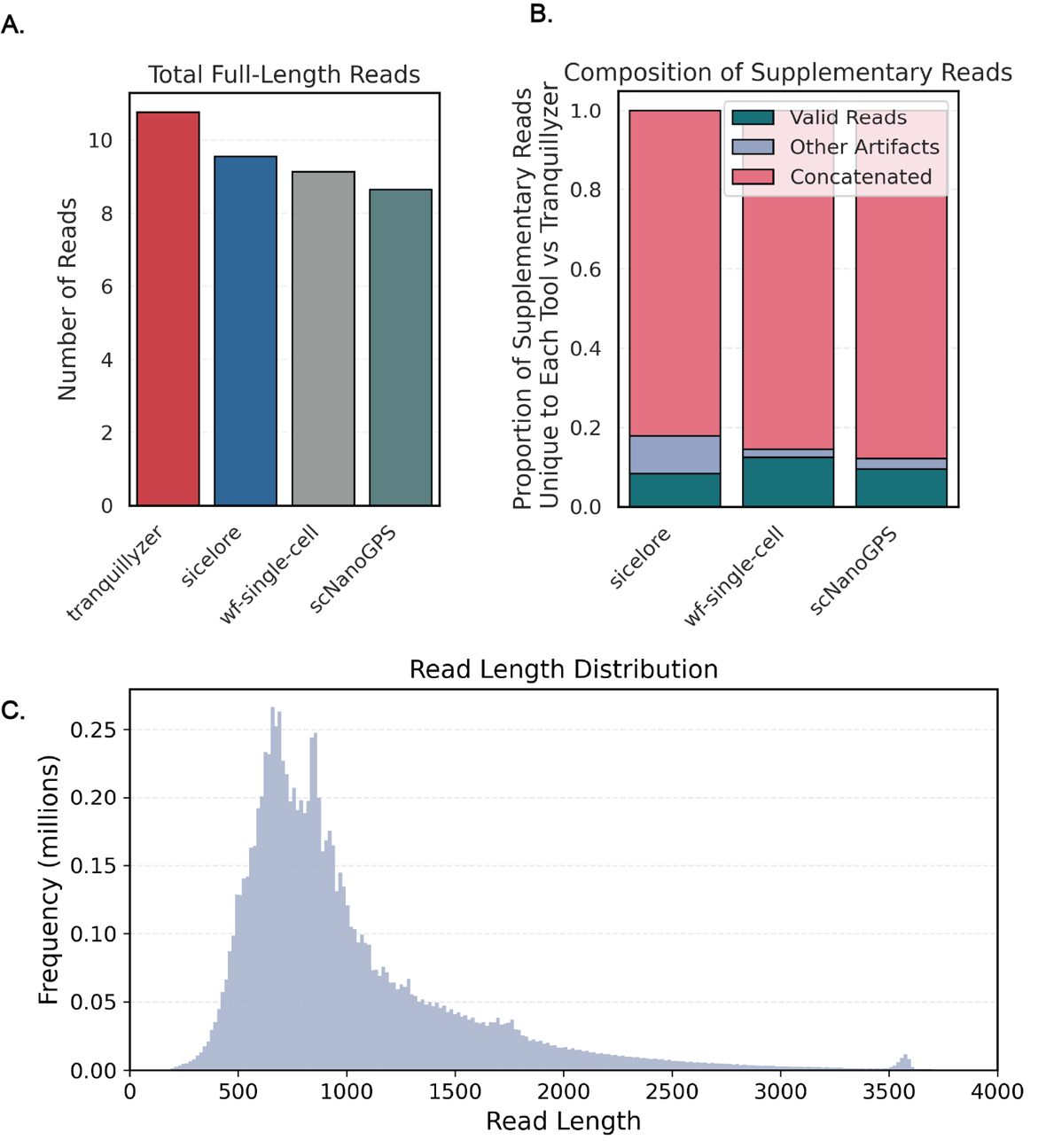

**Supplementary Figure S2: *Tranquillyzer* maximizes full-length molecule recovery while resolving artifactual complexity in real long-read single-cell transcriptomes.** (A) Read length distribution of the analyzed single-cell long-read dataset, showing a median read length of 855 bp with a broad range extending beyond 3,500 bp, including a small peak at ~3,600 bp indicative of possible concatenated artifacts. (B) Total number of full-length reads theoretically recoverable by each tool, calculated as the sum of valid single-fragment molecules and, for *Tranquillyzer*, the total sub-fragments that could be recovered by splitting structurally annotated concatenated reads. *Tranquillyzer* substantially outperforms other tools by leveraging its capacity to resolve multi-fragment molecules into their constituent parts. (C) Composition of supplementary reads uniquely identified by *Sicelore, wf-single-cell, and scNanoGPS*, benchmarked against *Tranquillyzer_CRF’s* structural annotations. Majority of supplementary reads from each tool (>80%) originated from reads *Tranquillyzer* had previously annotated as concatenated, indicating widespread misclassification of artifactual reads as valid by competing pipelines. These findings underscore *Tranquillyzer’s* robust capacity to resolve both canonical and complex multi-fragment molecules with high structural fidelity.
